# Positive Cardiac Inotrope, Omecamtiv Mecarbil, Activates Muscle Despite Suppressing the Myosin Working Stroke

**DOI:** 10.1101/298141

**Authors:** Michael S. Woody, Michael J. Greenberg, Bipasha Barua, Donald A. Winkelmann, Yale E. Goldman, E. Michael Ostap

## Abstract

Omecamtiv mecarbil (OM) is a positive cardiac inotrope in phase-3 clinical trials for treatment of heart failure. Although initially described as a direct myosin activator, subsequent studies are at odds with this description and do not explain OM-mediated increases in cardiac output. Single-molecule, biophysical experiments on cardiac myosin show that OM suppresses myosin’s working stroke and prolongs actomyosin attachment 5-fold, which explain inhibitory actions of the drug observed *in vitro*. Surprisingly, the increased myocardial force output in the presence of OM can be explained by cooperative thin filament activation by OM-inhibited myosin molecules. Selective suppression of myosin is an unanticipated route to muscle activation that may guide future development of therapeutic drugs.

## INTRODUCTION

Omecamtiv mecarbil (OM) is a positive cardiac inotropic agent that increases cardiac output in failing hearts, as measured by left ventricular fractional shortening and ejection fraction, in both animal models^1^ and humans^2–4^. OM was initially described as a direct binder and activator of β-cardiac myosin (MYH7), the molecular motor responsible for powering contraction^1^. The drug has generated great excitement, since a pharmaceutical that directly improves myosin activity holds the promise of avoiding side effects that can occur with drugs that target calcium signaling or other upstream regulators of contraction^5^.

Myosin activation by OM was proposed to be the result of an increase in the actin-activated rate of phosphate release (Figure 1, step 5) without changing the rate of ADP release (Figure 1, step 6)^1^. This kinetic modification is expected to increase the fraction of the ATPase cycle in which myosins are bound to actin in a force-bearing state (i.e., increasing the duty ratio), resulting in higher force production without affecting the rate of shortening^1^. Subsequent studies confirmed that OM increases the rate of phosphate release^6,7^; however, it was also shown that OM inhibits the velocity of actin gliding in the *in vitro* motility assay at all concentrations tested^6,8–10^ and reduces the rate of tension development and relaxation in myocytes at micromolar concentrations^1,9,11–13^. These phenomena, along with observations of decreased isometric force in fully activated cardiomyocytes,^11,14^ are inconsistent with the originally proposed model for OM-activation of myosin in muscle.

**Figure 1.**
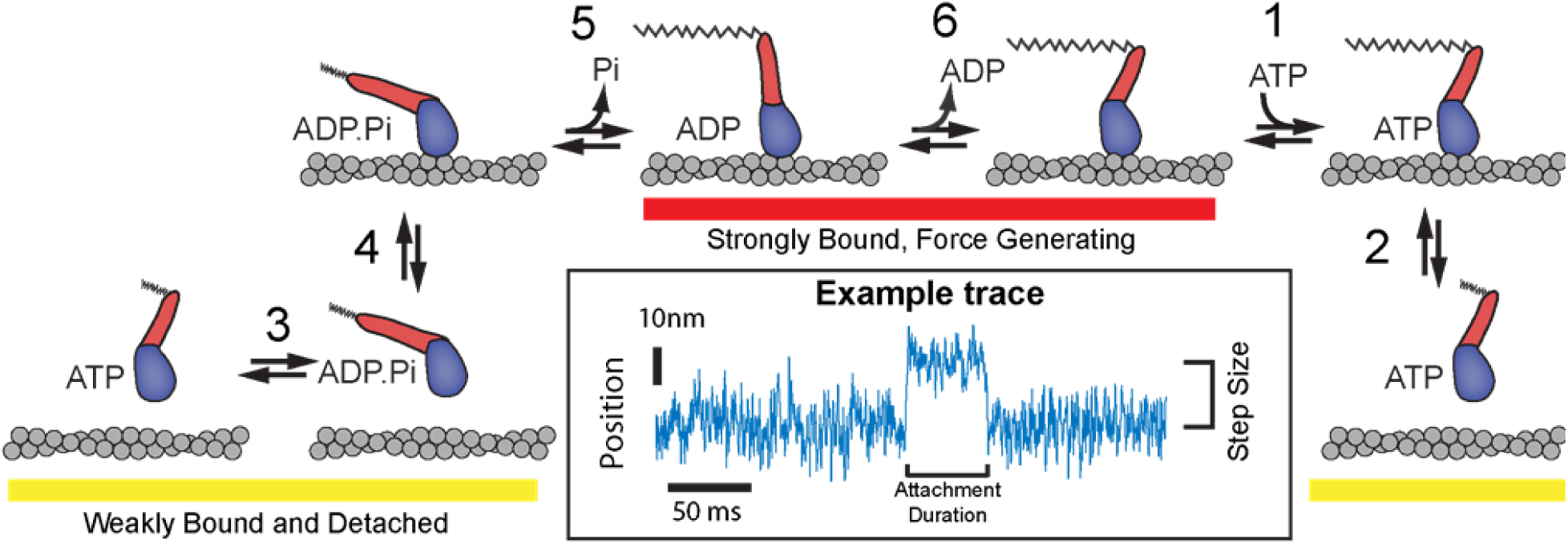
Biochemical cycle of cardiac myosin showing the key biochemical steps necessary for force production. Omecamtiv Mecarbil has been shown to increase the rate of phosphate release (step 5) and bias the ATP hydrolysis step (step 3) towards the post-hydrolysis M•ADP•P_i_ state, which is proposed to cause myosin to enter the strong binding states (red underline) more rapidly. Other biochemical steps have been previously shown through stop-flow biochemical experiments to be nearly unchanged by the presence of OM. Inset: Example optical trapping trace of the position (median filtered with 0.4 ms window) of an actin filament which occurs during one interaction with a single myosin molecule reproduced from Fig 2b. The step size and attachment duration of these interactions can be measured as shown.

To better understand the molecular mechanisms by which OM increases contractility, we utilized single molecule optical trapping to directly measure the effects of OM on the working stroke size and attachment lifetime of individual recombinant-expressed, human β-cardiac myosin molecules. The results of these experiments provide new information on the molecular mechanism of OM and how this drug can increase force production in hearts, while also inhibiting cardiomyocyte force production under high calcium and/or high OM concentrations and inhibiting *in vitro* actin gliding velocity. The results also point to a novel mode of muscle activation by the selective inhibition of a sub-population of myosin molecules.

## RESULTS

### OM Reduces the Size of Myosin’s Working Stroke

We kinetically and mechanically characterized the interaction of a recombinant, human β-cardiac heavy-meromyosin (HMM) construct with actin in the presence and absence of OM using single-molecule, optical trapping. In this assay, a single actin filament suspended between two optically-trapped polystyrene beads (known as an actin “dumbbell”) is brought into proximity with a single myosin molecule adsorbed to a pedestal bead which is anchored to the coverslip surface^15,16^. Single myosin molecules bind and displace the dumbbell when it is near the pedestal, and binding events are detected by analysis of the covariance of the bead position fluctuations^17,18^ (see Online Methods). Binding events as short as 12 ms can be detected, and the displacement of the dumbbell by the myosin working stroke is determined with sub-nanometer resolution (Figure 1, inset).

In the absence of OM, we observed directional displacements of the actin filament consistent with previous measurements (Figure 2a,b). Full length porcine β-cardiac myosin has been shown to have a working stroke composed of two sub-steps, where an initial displacement associated with the release of phosphate (4.7 nm) is followed by a second displacement associated with ADP release (1.9 nm)^19^. By averaging ensembles of single molecule interactions aligned at their beginnings and ends, we also resolved a two-step working stroke for human β-cardiac myosin at low MgATP concentrations (200 nM MgATP, Supplemental Figure 1). The initial displacement of 4.2 ± 0.4 nm is followed by a second displacement of 1.5 ± 0.3 nm, resulting in a 5.7 ± 0.3 nm total working stroke (Supplemental Figure 1). At a near-physiological MgATP concentration of 4 mM we observe a total working stroke of 5.4 ± 0.2 nm, but we do not fully resolve the second displacement due to the rapid binding of ATP and subsequent detachment of myosin (~500 s^-1^)^6^ immediately following the second displacement.

**Figure 2.**
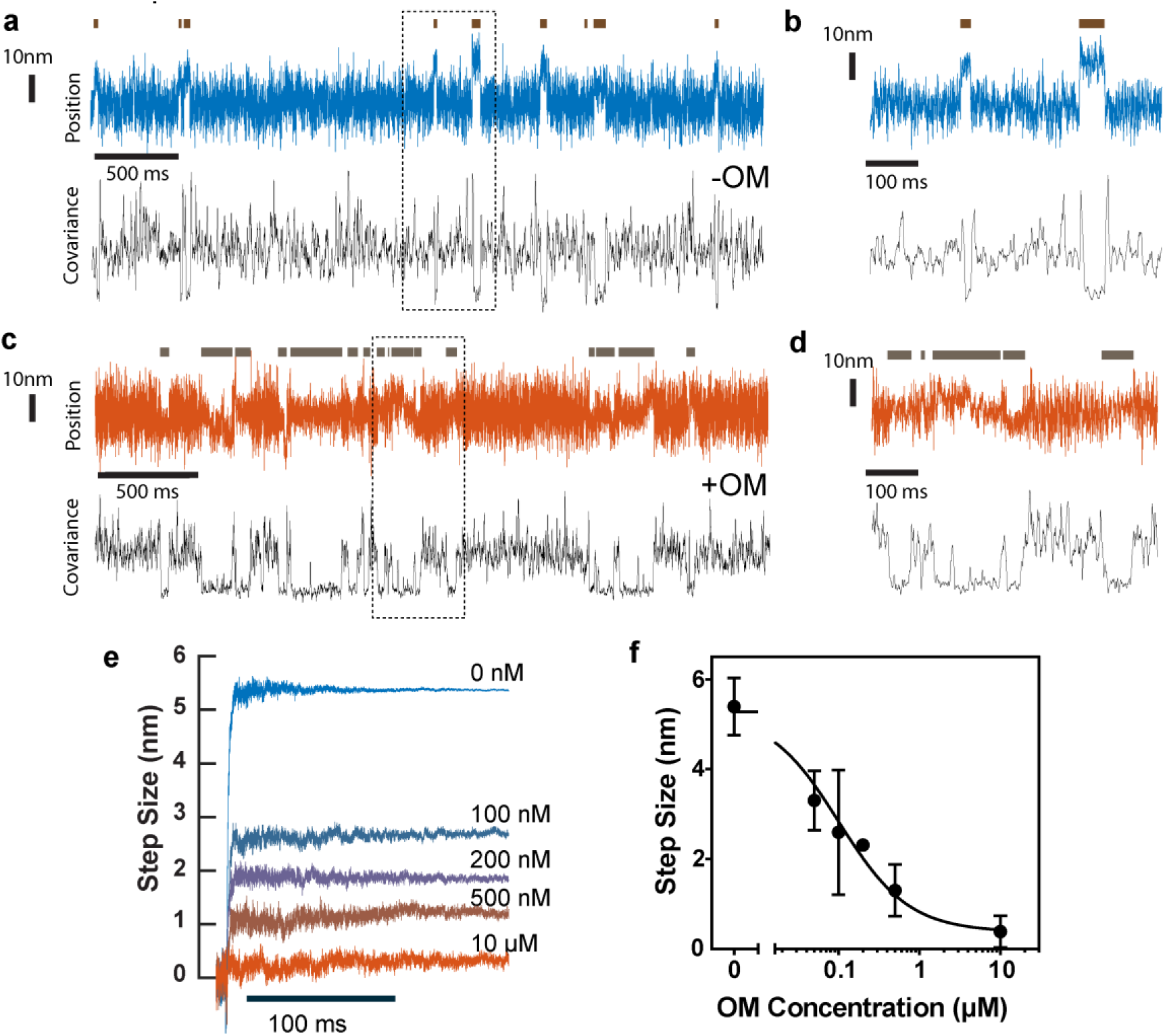
Step sizes of myosin at saturating MgATP (4 mM) and the effect of OM on the step size. **(a)** Example trace of the position of one bead during several interactions of cardiac myosin with actin in the absence of OM (blue). The covariance of the two beads’ positions is shown in black and was used to determine when a binding event occurred, as indicated by the dark horizontal lines above the position trace. **(b)** An expanded section of the data inside the dashed box in **(a)**, where two clear interactions can be visualized. **(c)** and **(d)** Example trace similar to that in **(a)** and **(b)**, but with 10 *μ*M OM present. The interactions are more difficult to distinguish in the position trace (red) but are clear from the covariance (black). Position traces in **(a)-(d)** are median filtered with a 0.4 ms window. **(e)** Binding events were synchronized at their starts and averaged forward in time to show the average step size observed in the presence of OM ranging from 0 to 10 *μ*M. Average step size decreases with increasing OM concentration. **(f)** The average observed step size was inhibited by OM in a dose dependent manner. Error bars give the standard deviation of the mean step sizes from each molecule observed. N-values are presented in Supplemental Table 1.

In the presence of a high OM concentration (10 μM), myosin attachments to actin were clearly resolved via a decrease in the covariance signal (Figure 2c, d). Strikingly, however, there was no clearly discernable working stroke observed (Figure 2e). The average observed step size was 0.4 ± 0.2 nm, and never exceeded 0.75 nm for any individual myosin molecule. The optical trap stiffness (0.06-0.08 pN/nm) was much lower than the stiffness of the myosin (0.5-2 pN/nm^20–22^) and nearly identical in the absence and presence of OM, so it is unlikely that the working stoke was suppressed by the small mechanical resistance imposed by the trap (~ 0.5 pN for a 5.5 nm displacement).

As the concentration of OM was varied from 0 to 10 μM, the mean working stroke was reduced from 5.4 ± 0.2 nm to 0.4 ± 0.2 nm in an OM concentration-dependent manner, with an EC_50_ of 101 ± 25 nM (Figure 2f).

We hypothesize that when OM is bound to myosin it completely inhibits the working stroke, and the probability of OM being bound to myosin during an interaction varies with OM concentration. We tested this hypothesis by assuming a model where the distributions of the individual step sizes were described by the linear combination of two Gaussian distributions, each with its own mean step size. We performed a MLE global fit across all observed OM concentrations in the software MEMLET^23^ to determine if the best-fit parameters from this model were consistent with our hypothesis (See Online Methods). The step sizes and widths of the two distributions were shared across all OM concentrations, while the relative proportion of each population could vary. The best-fit parameters to this model gave two populations with working strokes of 5.46 nm and 0.18 nm. The fraction of events with a mean working stroke of 5.46 nm decreased from 100% to 8.8% as the OM concentration increased from 0 to 10 μM with an apparent EC_50_ of 87.5 nM ± 31 nM (Supplemental Figures 2 and 3, Supplemental Table 2) which agrees with the EC_50_ for the average step size of 101 ± 25 nM. This result is consistent with a model in which myosin performs it full working stroke when OM is not bound but has virtually no net displacement when OM is bound, and the proportion of interactions that occur with OM is bound depends on the OM concentration.

**Figure 3.**
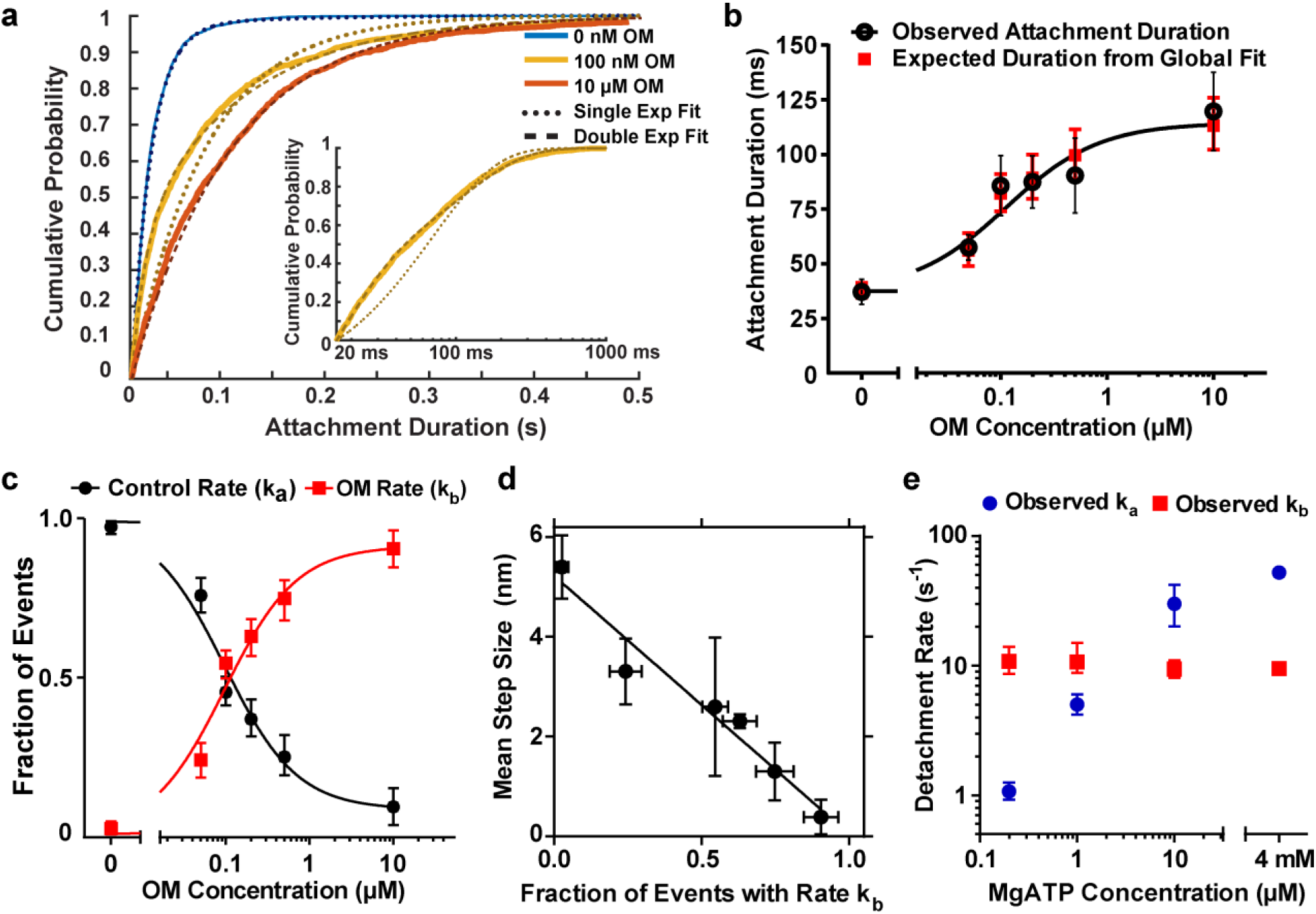
Actin attachment durations as a function of OM concentration. **(a)** Cumulative distributions of the actomyosin attachment durations (solid lines) at 4 mM MgATP. Without OM, the attachment durations are well described by a single exponential distribution (dotted blue line); however, at 100 nM and 10 *μ*M OM, double exponential distributions were required (yellow and red dashed lines). Inset: 100 nM OM durations on logarithmic x scale, highlighting the two phases of detachment. **(b)** Concentration-dependent effect of OM prolonging the mean observed attachment duration (black) at 4 mM MgATP. Black error bars show the standard deviation of the mean durations from each molecule. Red squares show the expected duration calculated from the global fit to durations. **(c)** The fraction of events which were found to detach at k_a_, (black), or at the OM-associated rate (k_b_, red) from the durations global fit as a function of OM concentration. **(d)** Observed step size linearly correlates with the fraction of events which detach at the OM-associated rate, k_b_. Vertical error bars are the standard deviation of the mean step from each molecule studied. **(e)** Detachment rates at 10 *μ*M OM as a function of MgATP concentration. Rate k_a_ (blue) was proportional to MgATP concentration at low ATP concentrations. Rate kb (red) was only observed in the presence of the drug and was independent of the ATP concentration at all concentrations studied. Unless otherwise noted, error bars from all panels show the 95% confidence intervals obtained via bootstrapping.

### Cycling Myosin Remains Attached to Actin Longer in the Presence of OM at Physiological ATP Concentrations

Actin bound durations were well-described by single exponential functions at all MgATP concentrations studied in the absence of OM. The rates were linearly related to MgATP at low concentrations (0.2 – 10 μM, Figure 3e), yielding an apparent second-order rate constant for MgATP binding and detachment (3.0 ± 0.14 μM^-1^s^-1^), which is consistent with biochemical measurements^6^. At saturating ATP (4 mM MgATP), the mean detachment rate of 47.7 s^-1^ (43.2-51.9 s^-1^ 95% CI, Figure 3a) is consistent with ADP release limiting the rate of actin detachment as predicted by biochemical experiments and shown previously^6,19,24^.

High concentrations of OM (10 μM) decreased the actin-detachment rate 5-fold to 9.4 s^-1^ (8.6 – 10.3 s^-1^ 95% CI), as calculated by a single-exponential fit. As the OM concentration varied from 0 to 10 μM, the mean observed attachment duration was prolonged with an EC_50_ of 114 ± 38 nM (Figure 3b, Supplemental Notes).

The durations of attachment with OM present were not well-described by single exponential distributions but were best fit by two-component exponential functions (Figure 3a, inset). At saturating ATP and at all OM concentrations studied, the rate of the faster phase (*k_a_*) is similar to the detachment rate of myosin in the absence of OM and the rate of the slow phase (*k_b_*) is similar to the detachment rate in the presence of 10 μM OM (Supplemental Figure 4, Supplemental Table 3). A global fit of a double exponential distribution was performed for the 0 – 10 μM OM datasets simultaneously with rates *k_a_* and *k_b_* being shared between the data sets. Only the relative proportion of events which dissociated at each rate was allowed to vary between OM concentrations^23^. This procedure yielded two common rates similar to those observed at the 0 μM and 10 μM OM concentrations (*k_a_* = 52 s^-1^ and *k_b_* = 9.4 s^-1^), and the relative fraction of events dissociating at rate *k_b_* increased monotonically with increasing OM concentration with an EC50 of 114 ± 28 nM (Figure 3c). This supports the idea that the concentration of OM affects the likelihood that myosin will be bound to OM when it binds to actin, and that when OM is bound, the rate of detachment of myosin is changed from *k_a_* to *k_b_*.

**Figure 4.**
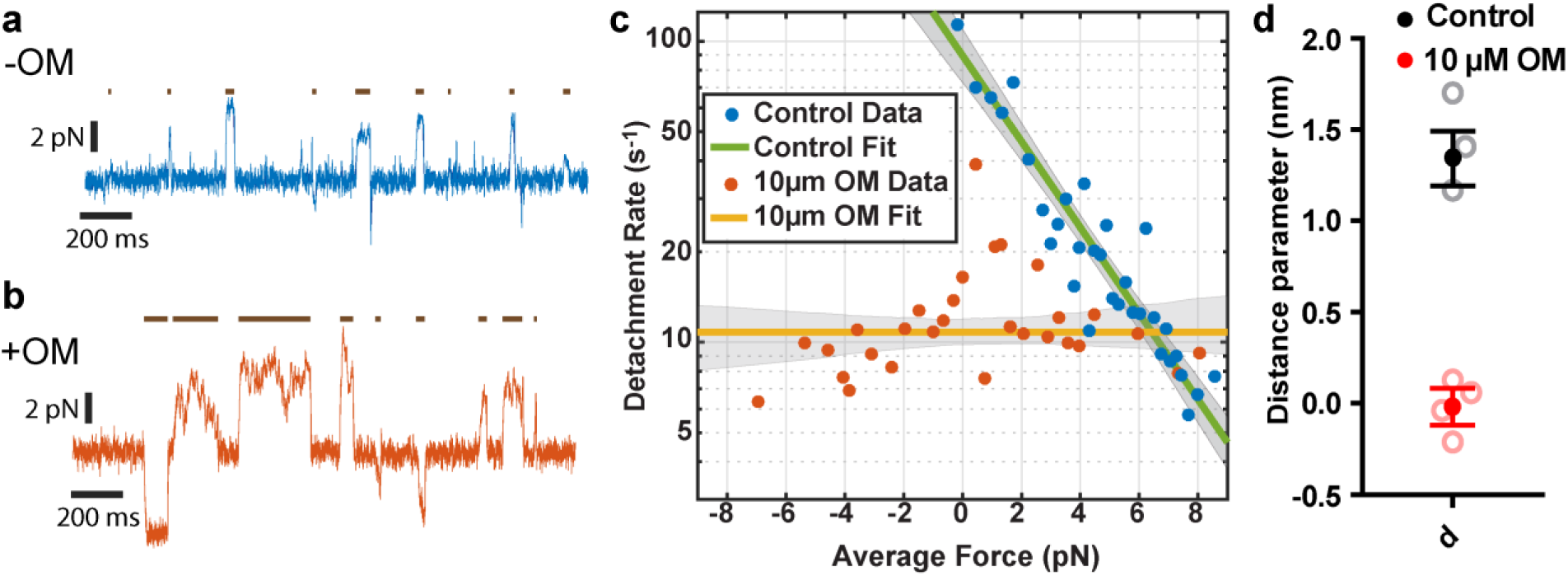
Force dependence of detachment is eliminated by OM. **(a)** and **(b)** Example traces of force on the motor bead (blue) with the trap’s isometric feedback system engaged. Events, as detected by covariance, are indicated by dark lines above the force traces. **(a)** In the absence of OM (blue), forces are predominantly in the positive direction (myosin is under resisting load) **(b)** In the presence of 10 *μ*M OM (red), forces are generated in both directions. **(c)** The observed detachment rates in the absence of 0 OM (blue) and at 10 *μ*M OM (red) as a function of applied load are shown as circles at the average force and rate of 20 events sorted by force. Green and yellow lines show the detachment rate calculated from Equation 1 and parameters from the MLE fit in the absence of and presence of 10 *μ*M OM, respectively, with 95% confidence intervals shown as shaded grey areas. **(d)** Distance parameter estimates for the control (no OM) and 10 *μ*M OM data from all observed molecules (closed circles) are shown with 95% confidence intervals from bootstrapping and the estimated distance parameters from individual molecules (open circles).

The proportions of events detaching at the OM-associated rate, *k_b_*, show a linear relationship with the observed step sizes (Figure 3d). The observed step decreased as more events detached at rate *k_b_*. This relationship supports a model in which both the step size is suppressed and the detachment rate is changed when OM is bound to myosin.

### Dissociation of Actively Cycling, OM-Bound Myosin is Independent of MgATP Concentration

At low ATP concentrations (≤ 10 μM), where the actomyosin dissociation rate is dependent on the ATP concentration, the kinetics of actomyosin dissociation with OM present were again best described by a two-component exponential distribution (Supplemental Figure 5). One of the observed rates (*k*_b_) was near 10 s^-1^ for all MgATP concentrations studied (Figure 3e, red squares). The other rate (*k*_a_) varied with ATP concentration and was consistent with the previously measured second order rate constant of 3.0 μM ^-1^ s^-1^ for ATP binding to actomyosin. At MgATP concentrations ≤1 μM MgATP, the ATP-dependent rate of detachment was *slower* than the OM-associated 10 s^-1^ detachment rate, while at MgATP concentrations ≥10 μM, ATP-induced dissociation was *faster* than 10 s^-1^ (Figure 3e). Detachments with rates *k*_a_ and *k*_b_ were observed under the same conditions in a single experiment from the same molecule. This result indicates that after OM-bound M•ADP•Pi binds to actin and enters a strongly bound state (step 5; Figure 1), its subsequent actin dissociation is not through ATP binding to AM (step 1; Figure 1). Additionally, the dissociation at rate *k*_b_ is not due to OM-induced dissociation of the M or M•ADP states from actin, as stopped-flow biochemical experiments indicate that these dissociation rates are substantially slower than *k_b_* and not affected by OM (Supplemental Notes, Supplemental Figure 6, Supplemental Table 4). Taken together these results indicate that cycling, OM-bound myosin does not pass through the canonical rigor state that requires ATP binding for detachment, but rather follows a non-canonical pathway.

**Figure 5.**
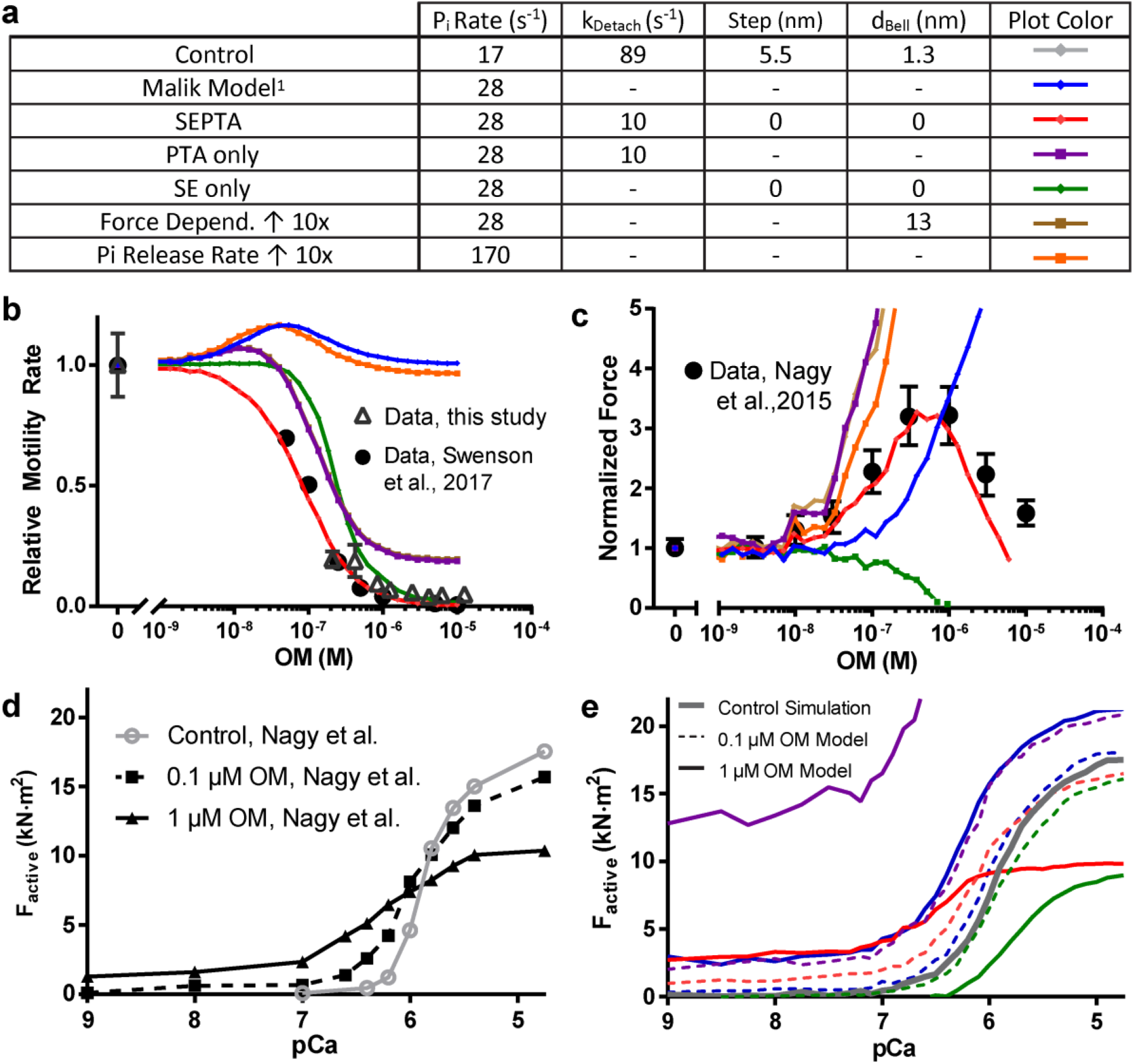
Simulations of OM’s effect on gliding velocity and isometric force. (**a)** Summary of parameters used in models in panels **(b)**-**(e)**. **(b)** Comparison of gliding filament velocity for the protein used in this study and from Swenson et al^*9*^. to simulated data from the various model parameter sets (colored lines as in **(a)**). Only the SEPTA Model (Stroke Eliminated, Prolonged Time of Attachment), with parameters from the single molecule measurements, fully accounts for the observed marked decrease in vitro velocity as a function of OM concentration. The curve for 10X Increased Force Dependence is hidden behind the curve for PTA only (Prolonged Time of Attachment only, purple). Motility error bars are standard deviations of velocities from individual filaments. **(c)** Comparison of simulated, normalized, isometric forces at an intermediate calcium concentration (15% activation, lines colored as in **(a)**) and results from Nagy et. al^*11*^ who reported a bell-shaped force response of permeabilized myocardial trabeculae as a function of OM (black circles). Only the SEPTA model (red) shows a similar biphasic shape. **(d)** Active force data reproduced from Nagy et. al. as a function of pCa (-log [Ca^2+^]) at 0, 100 nM and 1 *μ*M OM. **(e)** Simulated isometric force at 0 (grey line), 100 nM (dashed lines, colors as in **(a)**), and 10 μM OM (solid lines) for comparison with the experimental data in **(d)**. Only the SEPTA model (red dashed and solid curves) recapitulates the leftward shift in the pCa-tension curve (calcium sensitization) and decreased force production at fully activating [Ca^2+^].

**Figure 6.**
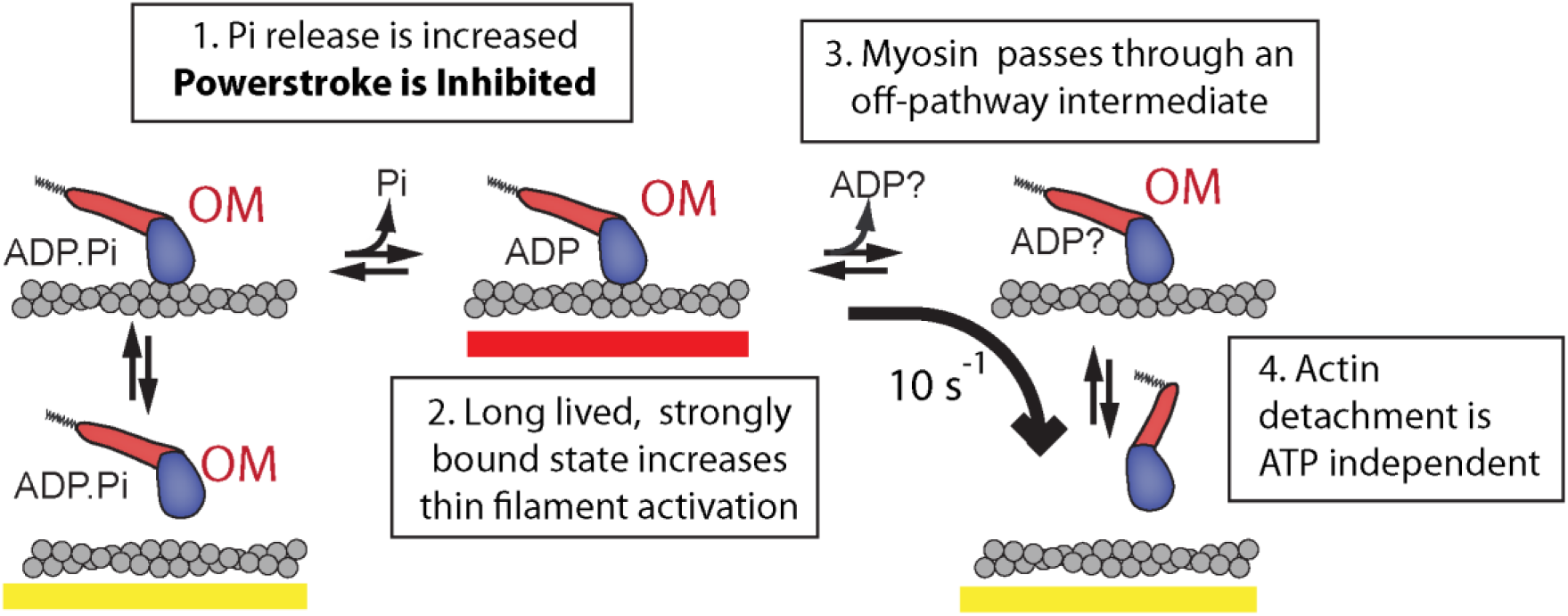
Model of OM’s effect on cardiac myosin. 1) OM increases the rate of entry into strong binding as previously measured by phosphate release rates, but the force generating power stroke is inhibited. 2) Myosin remains strongly bound to actin, contributing to increased thin filament activation at intermediate calcium concentrations. 3) OM disrupts the typical pathway of myosin, causing it to pass through an ADP or apo state with its lever arm still in the pre-power stroke position. 4) Myosin detaches from actin without needing to bind ATP. ATP binding must occur before the cycle can start again.

### Force Dependence of Detachment Rate is Eliminated by OM

To probe the force dependence of the attachment lifetime of cardiac myosin, we used an isometric feedback technique^18^ previously used to determine the force dependence of various myosin isoforms^16,19,25^. This technique applies a load onto the myosin molecule by varying the position of one of the optical traps to maintain a constant position of the bead in the opposite trap. This acts to maintain the actin filament (which is inside the feedback loop) in a nearly constant position, allowing myosin to exert an isometric force while it interacts with actin. The value of the applied load is determined by the distribution of myosin binding positions along the actin and the magnitude of the power stroke. In the absence of OM, myosin was able to maintain isometric forces ranging from −1 pN to +7 pN (Figure 4a), while in the presence of 10 μM OM, the observed forces ranged between −8 pN to +8 pN (Figure 4b). There was no apparent bias to the direction of the applied forces with OM present, even within a single molecule’s interactions with a single actin filament. The force distribution in the presence of OM is likely due to stochastic binding along the actin filament to Brownian Motion unbiased by a working stroke, supporting the result given earlier (Figure 2) that OM suppresses the working stroke.

The force-dependence of the actin-detachment rate can be quantified by fitting the Bell equation^26^ to the attachment durations without binning using Maximum Likelihood Estimation^23^:

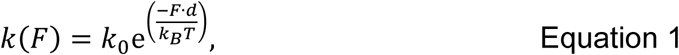

where *k_0_* is the rate of detachment in the absence of applied load, *F* is the applied force, d is the distance to the force-dependent transition state and a measure of the degree of force dependence, *k*_B_is Boltzmann’s constant, and *T* is temperature. Binned detachment rates as a function of force and the results of fitting Eq. 1 to the data are shown in Figure 4c. In the absence of OM, we found that the distance parameter (*d*) was 1.3 nm (1.19-1.49 nm 95% CI), with *k_0_* = 89 s^-1^ (72.9-109 s-1 95% CI), consistent with previous measurements for porcine β-cardiac myosin^19,24^. However, in the presence of 10 μM OM, there was no appreciable change in the detachment rate as a function of load (Figure 4d). A log-likelihood-ratio statistical test^23^ also indicates that fitting the Bell model to the data is not justified over a force-independent single exponential distribution (p = 0.62).

### Calcium Sensitization and Motility Inhibition by OM is Explained by Suppressed Step Size and Prolonged Attachment Duration

We carried out simulations to determine whether the results we observed in the optical trapping assay could explain the effect of OM on the actin gliding rate in the *in vitro* motility assay^6,8–10^and on the calcium-dependence of contractile function of cardiomyocytes^11,14^. We used a simple, two-state model of the actomyosin cycle along with cooperative activation of the thin filament as simulated by Walcott et al. ^27,28^ (See Online Methods). This model was used to test whether the effects of OM on attachment rate, detachment rate, step size, and force dependence would alter gliding velocity and isometric force as a function of OM and calcium. We utilized parameter values from the literature^6,20,28,29^ combined with the detachment rate, step size, and force dependence measured in our experiments in the presence and absence of OM (Supplemental Table 5). We tested six possible models of OM’s effects (Figure 5a), including one where only the phosphate release rate was increased (Malik Model). All other models also included this increase in phosphate release rate, but varied in the size of the step, the detachment rate, and/or the force dependence of detachment rate. The model utilizing the parameters measured in this work for step size, force dependence, and attachment duration is referred to as the SEPTA Model (<u>S</u>tep <u>E</u>liminated, <u>P</u>rolonged <u>T</u>ime of <u>A</u>ttachment).

Simulations of unloaded velocity, using an OM binding affinity of 100 nM, showed that the SEPTA model (Figure 5b, red) which includes a decreased step size and decreased detachment rate was able to fully account for the drastic reduction in actin gliding velocity we and others^8–10^ observed *in vitro* (Figure 5b, black circles). It is also possible to simulate a drastic reduction in the motility rate by decreasing the working stroke size without changing the detachment rate (Figure 5b, SE model, green). The Malik Model, the previously proposed mechanism of action for OM^1^, did not decrease the velocity (Figure 5b, Malik, blue). It had previously been suggested that the slowed motility could be due to increased force dependence of myosin detachment in the presence of the drug, but even a 10-fold increase in the distance parameter, *d*, corresponding to a greatly increased force sensitivity of detachment could not reproduce the experimental motility assay data (Figure 5b, brown). Increasing *d* 10-fold effectively reduces the mean detachment rate to ~10 s^-1^ causing the outcome of this model to match the results of the PTA Model where only the detachment rate was changed (Figure 5b, purple).

Experimental measurements using isolated and permeabilized cardiomyocytes show a biphasic effect of the OM concentration on isometric force production, with force enhancement peaking at intermediate calcium concentrations^11,14^. The SEPTA model reproduced this biphasic force response in simulations (Figure 5c, red). Simulations using other parameter variations were able to predict an increase in force production with increasing OM, but only when the working stroke was inhibited did the force decrease at high simulated OM concentrations (SEPTA, SE Models). SEPTA also was the only one of the six combinations tested that reproduced the inhibition of force by OM at high calcium concentrations (pCa 4.75), and the increased calcium sensitivity reported by Nagy et. al and others^9,11,13,14^, who demonstrated a leftward shift in the pCa-tension curve (Figure 5d-e, red). Our simulations of myocyte isometric force used a dissociation constant for OM of 1.2 μM in contrast with 100 nM found in our optical trapping data and used for the *in vitro* motility simulations. This 1.2 μM K_D_ is consistent with the observed effect of the drug in muscle fibers as discussed in the Supplemental Notes.

## DISCUSSION

OM is not a direct activator of myosin, and its inhibitory effects on motility seem counterintuitive for a drug that improves myocardial function in heart failure. Our single molecule results provide a revised mechanism of action of this interesting pharmaceutical that explains results from a variety of biochemical, structural, and physiological experiments.

Our optical trapping studies indicated that OM binding to myosin has two main effects. First, it inhibits the size of the working stroke more than 10-fold, from 5.4 nm to less than 0.4 nm. Secondly, actomyosin attachment duration is prolonged at physiological ATP concentrations by 5-fold, and detachment becomes independent of both ATP concentration and force applied to the myosin. The OM effect on both the working stroke and attachment duration occur with a *K_i_* of ~100 nM. A simple model predicts that prolonged attachments lead to increased force production in muscle due to thin-filament activation (see below), while the inhibited step size explains the dramatic slowing of gliding velocity *in vitro* and decreased force production in cardiomyocytes observed under either high OM concentrations or at all OM concentrations for saturating calcium concentrations.

Our discovery of the OM-induced decrease of the myosin working stroke size clarifies the findings of previous studies. Biochemical and structural studies demonstrated that OM stabilizes the pre-powerstroke ADP•P_i_ state^6^,^30^, and transient-time-resolved distance measurements utilizing fluorescence resonance energy transfer suggest that the rate of a conformational change that correlates with myosin’s working stroke is drastically reduced by OM^7^, leading to the possibility that myosin detaches from actin before the power stroke occurs. Additionally, molecular dynamics studies of cardiac myosin bound to OM suggest that movement of the myosin lever arm relative to the myosin motor domain may be inhibited by OM^31^. Recently reported *in-situ* studies also have shown evidence that the stroke may be reduced by OM in muscle fibers^32^.

OM had been proposed not to affect the actomyosin detachment rate during cycling^1,6^, which is at odds with our findings. Previous conclusions were based on the biochemical measurement of ADP release (the step that limits the rate of actin detachment at physiological ATP) from an AM•ADP complex formed by adding ADP to the rigor, AM complex ^1,6,9^ Our results indicate that the step that limits actin detachment is not accessible by simply adding ADP, but occurs earlier in the cycle and may not be on the conventional ATPase pathway (Figure 6). Interestingly, a single-turnover, stopped-flow study showed the rate of ADP release was slowed by OM when myosin proceeded through its cycle from the M•ADP•P_i_ state^9^. However, the authors of this study concluded that the actin-bound, ADP-isomerization step (AM’•ADP −> AM•ADP) found on the canonical ATPase pathway was affected by OM. This interpretation is inconsistent with data presented here (Figure 3e), since their model would include a normal working stroke displacement and an actin detachment rate that would be slowed by low MgATP concentrations.

In contrast, our experiments suggest that myosin detaches from actin from an ADP bound or apo state off the typical pathway, since we observe that myosin does not undergo a working stroke while attached to actin. This finding is consistent with the lack of force dependence of actin detachment rate in the presence of OM (Figure 4). Forces that resist the myosin working stroke have been shown to slow the ADP isomerization or ADP release step^19,24^. As the working stroke does not occur in the presence of OM, the post-stroke state preceding force-sensitive ADP release is never populated, and the kinetic step that limits detachment is thus not force dependent.

Our SEPTA (<u>S</u>troke <u>E</u>liminated, <u>P</u>rolonged <u>T</u>ime of <u>A</u>ttachment) model reproduced the biphasic effect of isometric force production on OM concentration in cardiomyocytes (Figure 5). Models that included increased phosphate release rates and and/or prolonged actin attachment resulted in monotonic increases in force with increasing OM (Figure 5c). The force increase is expected, since increasing the phosphate release rate and decreasing the actin detachment rate lead to a higher duty ratio, with more myosins dwelling in the strongly-bound states. However, increasing phosphate release (Figure 5, Malik Model) and/or prolonging the time attached (Figure 5, PTA only) alone does not account for the experimentally observed decreases in force at high OM concentrations or at fully activating calcium concentrations (Figure 5c,e). An OM-induced decrease in the working stroke does account for these effects. Although not modeled in our simulations, OM’s proposed effect on thick filament activation also could result in monotonically increasing force production with OM concentration.^14^ While thick filament activation alone cannot explain OM’s biphasic effect on force production, it may explain the observed decrease in the Hill coefficient for the isometric force pCa curves which is not reproduced in our simulations.

Our simulations suggest that the experimentally observed increase in cardiomyocyte force production in the presence of sub-micromolar OM concentrations are due to prolonged actin attachment increasing thin filament activation. OM-bound myosins do not produce a power stroke and cannot generate force, so experimentally observed increases in force must be due to the increased recruitment of drug-free myosin. OM-bound myosin activates the thin-filament regulatory system allowing the cooperative binding of fully functioning myosin^33^. A similar effect on thin filament activation has previously been shown in low ATP conditions, where the actin-attachment duration of myosin to the thin filament is increased^34,35^. At high concentrations of the drug, when a majority of myosin heads are bound to OM, force is inhibited at all calcium concentrations, since OM-bound myosin does not perform its working stroke. A prediction of the SEPTA model, that is seen experimentally^11,13,14^, is that OM does not increase force production at calcium concentrations that fully activate the thin filament, and increasing OM concentrations inhibit force. The maximal force at intermediate calcium (15% activation) is developed when approximately 30% of myosin heads are bound to OM in the conditions of our simulation (Supplementary Figure 7).

The observed effects on working stroke and attachment duration occur at OM concentrations lower than the therapeutic plasma concentration of OM in patients of approximately 200-600 nM^2,4^, making these observations relevant to therapeutic applications of the drug (See Supplemental Notes).

## CONCLUSIONS

Our single molecule, optical trapping experiments have revealed that at concentrations relevant to therapeutic treatment of patients, OM acts to suppress cardiac myosin’s working stroke and to prolong the time of actin attachment. These effects intuitively seem at odds with the positive results of the clinical trials which showed increased cardiac output from patients treated with the drug. However, these findings can account for the previously unexplained inhibition of the gliding velocity *in vitro* and force production in cardiomyocytes under saturating calcium and/or high OM concentrations. The previous hypothesis that motility was inhibited by increased force sensitivity of the myosin detachment rate has been shown to be unlikely, both by direct measurement of the force dependence (Figure 4) and our modeling results (Figure 5). Our simulations show that the increased attachment lifetime of a small population of non-force generating myosin molecules is sufficient to explain the calcium sensitization effect observed in cardiomyocytes and is the likely explanation for the physiological benefits seen in clinical trials. The concentration of OM in patients has been shown clinically to be crucial, as too much of the drug slows contraction to the point where arrhythmias or other adverse reactions may occur. This new knowledge helps to resolve the apparent inconsistencies observed in the wide variety of experiments and applications that have used this promising drug. In addition, it suggests a new approach for muscle-targeted drug development, showing that drugs which selectively inhibit a population of myosin may be able to cause increased force development *in vivo*.

## METHODS

### Protein Purification, *in vitro* Gliding Assays, and OM Source

Human β-cardiac myosin was purified and actin gliding assays were performed as described previously^10^. OM was obtained from Selleck Chemical (S2623) and a 10mM stock solution was prepared in DMSO and stored at −80 C.

### Optical Trapping Assay

Flow cell chambers were constructed as previously described using double-sided tape and vacuum grease^19^. The surface of the coverslip was coated with a 0.1% nitrocellulose solution (EMS) mixed with 2.5 μm diameter silica pedestal beads (Polysciences). The main assay buffer (AB) contained 25 mM KCl, 60 mM MOPS, 1 mMDTT, 1 mM MgCl_2_, and 1 mM EDTA. Cardiac myosin in myosin buffer (AB with 300 mM KCl) was incubated in the 20-30 μL chamber for 30 s to non-specially adhere to the nitrocellulose coated surface, before being washed out with additional myosin buffer. The concentration of myosin, ranging from 0.02 to 0.1 mg/mL, was adjusted each day so that only 1 out of 5-10 locations tested showed interaction with the actin filament. The chamber was blocked with 2 incubations of 1 mg/mL BSA each lasting 3 minutes. The experimental solution was added to the flow cell containing MgATP, either 0.1% DMSO for control experiments or OM (in DMSO with a final DMSO concentration of 0.1% for all conditions), 0.1-0.2 nM actin filaments composed of 10% biotin actin (Cytoskeleton) and 90% unlabeled rabbit skeletal actin prepared as previously described^36^ and stabilized by rhodamine labeled phalloidin (Fisher) and an oxygen scavenging system of approximately 3 mg/mL glucose, and glucose oxidase catalase. For ATP concentrations ≤ 10 μM, the concentration of a freshly made 1 mM MgATP stock solution was verified spectroscopically each day. Polystyrene beads with a diameter of 500 nm (Polysciences) were prepared by incubating approximately 0.40 ng of beads with 10 μL of 5 mg/mL neutravidin solution (Thermo) in water overnight at room temperature before washing 4 times with AB via centrifugation. Four microliters of a solution containing approximately 5 ng/mL neutravidin-coated beads, MgATP, and either 0.1% DMSO or OM was added to the chamber before it was sealed with vacuum grease.

Experiments were performed on a dual-beam optical trapping setup^37^ which utilizes a 1064 nm laser for trapping and direct force detection using quadrant photodiodes (JQ-50P, Electro Optical Components Inc.) and a custom-built amplifier. The two beams were produced and controlled by splitting the laser by polarization and passing each beam through a 1-D electro-optical deflector (LTA4-Crystal, Conoptics), which could deflect the beam based on input from a high voltage source (Conoptics, Model 420 Amplifier). Data acquisition, feedback calculations, and beam position control output utilized a LabVIEW Multi-function I/O device with built-in FPGA (PXI-7851) and custom built virtual instruments (LabVIEW). Data acquisition and digital feedback calculations occurred at 250 kHz.

The microscope utilized a Nikon Plan Apo 60x water immersion objective (NA 1.2) and a Nikon HNA Oil condenser lens (NA 1.4). One bead was trapped in each of the two 1064nm beams, with a trap stiffness of 0.06 −0.08 pN/nm, calculated via the power spectrum of the beads’ positions^38^. An actin filament 5-10 μm long was attached at each end to the two beads. The position of one bead was adjusted (using a servo-controlled mirror that was positioned conjugate to the back focal of the objective) to stretch the actin filament under 4-6 pN of pretension, reducing the effects of the non-linear compliance of the bead-actin linkage. Pedestal beads were tested by moving the actin filament close and observing whether any reductions in variance of the measured force signals occurred. When interactions were detected, a piezo-electric stage (Mad City Labs) was used to fine tune the position to maximize the number of interactions per second. A feedback system using an image of the pedestal bead and the nano-positioning stage was used to stabilize the stage position as previously described.^37,39^ For a given molecule, the position of the stage was slightly shifted by 6-12 nm between some data acquisition traces to reduce inhomogeneity of the accessibility of the actin attachment zones^21^.

The isometric feedback experiments were conducted as previously described ^16,19,25,40^, but with a digital feedback loop and EODs to steer the beam position. Briefly, a feedback loop held the position of one of the beads (referred to as the transducer) constant by modulating the position of the other trap, known as the motor trap. Since the actin filament is between the two beads and is inside the feedback loop, its position was maintained continuously, allowing the myosin to develop isometric force during its interaction with actin. The excursion of the motor trap was limited to 100-125 nm (corresponding to approximately 6-10 pN) to prevent entering the non-linear force regime due to the bead being pulled too far from the trap center. The response time of the feedback loop during myosin interactions was approximately 15-20 ms.

### Optical Trap Data Analysis

Data were analyzed as previously described^16,19,25^, by calculating the covariance of the two beads with a window of 8-15 ms. The covariance signal from a 30 s recording was fit to a double gaussian distribution, with the mean of the high covariance value peak representing when myosin was detached and the lower covariance value indicating attachment to actin. Events were selected by determining when the covariance signal passed from the detached value to the attached value and back to the detached level. The time of the binding event start and end were refined by examining the point when the covariance crossed a threshold calculated to minimize the overlap between the distributions of bound and unbound covariance values^16,25^. The duration of the event was calculated from these values of the start and end of each event. The total step size of the event was found by averaging 1 ms of the actin displacement of both beads 4-10 ms prior to dissociation and subtracting the baseline displacement of the actin position calculated by the average of a 1 ms window 4 ms after the detected end of the event. Depending on experimental conditions (dumbbell length, stiffness of bead-actin linkages, etc) events shorter than 12-24 ms could not be reliably detected and were eliminated from the analysis. This deadtime was determined by taking the size of the covariance window and multiplying it by a factor of 2. Ensemble averaging was performed as previously described, and were weighted such that each molecule contributed equally to the average.^16,19,25,41^

Events from data acquired utilizing isometric feedback were also analyzed by examining the covariance signal, which clearly decreased during actomyosin interactions. While the feedback loop was engaged, we also observed transient (<5 ms) rises in the covariance signal in the middle of interactions that were plainly not associated with detachment but were more likely due to a small conformation change in the myosin which causes both beads to move together simultaneously. These transients in the covariance signal were recorded as detachment events by the initial analysis, but a second pass through the data eliminated these detachments by removing detected unbound events which both lasted less than 2 times the covariance calculation window and were not associated with a return of the motor bead force to the baseline. The average force during an interaction was calculated by averaging the force on the motor bead starting 2 ms after detected attachment through 2 ms before detachment. The baseline force on the motor bead 4 ms after detachment was subtracted from this average force.

### Attachment Duration and Step Size Parameter Estimation

Detachment rates and mean step sizes were calculated using MEMLET^23^, a MATLAB based program which utilized maximum likelihood estimation to perform parameter estimation without the need for binning. For each set of conditions, parameter estimation was performed on data combined from multiple molecules (estimates from individual molecules yielded similar results to the combined datasets). Because the number of recorded events from each molecule varied due to non-biologically relevant experimental conditions (age of chamber, precise positioning of the actin filament, number of data traces recorded by the user), for each condition, the data was weighted such that the data from each molecule counted equally in the parameter estimation process. The multidimensional fitting capability of MEMLET was used to include weights for each duration or step size, and each weight included as an exponential factor acting on the total probability density function (PDF) describing the distribution before the log of the PDF was minimized as previously described^23^. The weights were calculated such that the sum of all weights was equal to the number of total points being fit, so that the estimated log-likelihood would be consistent with unweighted fitting results. Duration data were fit to either single exponential or double exponential PDF with weights, while step size data was fit to a weighted double Gaussian distribution, as in MEMLET (v 1.3).

Global fitting was performed on the step size distribution data to determine the parameters for a two-component gaussian model. In this weighted global fit, data from each OM concentration contributed equally to the parameter estimation, and data from each molecule contributed equally within a given OM concentration. Unweighted fitting produced similar results to this weighted approach. In the global fit, the means and standard deviations were shared between all datasets, but the relative amplitude of the two components could vary. Using a double Gaussian distribution to fit each condition individually was not statistically justified over a single component, as measured by the log-likelihood ratio test. Simulations showed however that this would be expected even if two populations of events were present with the distributions obtained via global fitting. This is because the relative size of the standard deviations of the step size distributions (approximately 7-8 nm) is large compared to the expected difference between the mean step sizes (~0 nm and ~5.5 nm), and even several thousand simulated event measurements was not enough for 2 components to be statistically justified.

Weighting of molecules and conditions for the event duration global fit was performed as described above for the step size global fit.

### Fitted Parameter Calculations and Reported Uncertainties

Calculations of the effective concentration leading to 50% of the observed effect (EC_50_) were performed in Prism v 7.03 (GraphPad Software). Three parameters were used in the model (Min, Max, EC_50_), and the standard error of mean was reported for EC_50_. Uncertainties in step size measurements are reported as standard errors of the mean, calculated from the step size distributions. Uncertainties for rates and distance parameters are given as 95% confidence intervals calculated via 200-500 rounds of bootstrapping^23^, corresponding with approximately with twice the expected standard error of the mean.

### Stopped Flow Experiments

Transient kinetic experiments were performed using pyrene labeled actin filaments as previously described^42^. Porcine cardiac myosin was prepared from cryoground pig heart ventricles (PelFreeze) as described by Kielley and Bradley^43^ and was then digested by chymotrypsin to isolate the S1 fragment.^44^ Experiments were performed using the same motility buffer as used in the trapping experiments at room temperature (20 °C). All solutions contained a total of 0.1% DMSO. For AM•ADP dissociation experiments, 1 μM myosin S1 was incubated with 2 mM MgADP and 1 μM pyrene labeled F-actin before being rapidly mixed with a solution containing 76 μM unlabeled F-actin, 2 mM MgADP, and 0.02 units/mL apyrase. The pyrene fluorescence transient was fit to a single exponential function plus a linear function (to account for drift over the long acquisition time). The rigor myosin dissociation experiments were conducted with similar solutions with the absence of MgADP and apyrase.

### Simulations

A Monte Carlo simulation using a modified Gillespie method^45^ was utilized to relate the single molecule biochemical and mechanical measurements to the previously described results from in-vitro motility assays^9,10^ and permeabilized cardiomyocyte assays^11,14^. To keep the model simple, we only included two states of the myosin, bound and unbound to actin, and ATP hydrolysis was ignored. A single thin-filament from half a sarcomere was modeled, where 75 myosin molecules spaced 14.3nm apart (coming from 3 thick filaments) could interact continuously with the infinitely stiff thin filament. Cooperative activation arising from strong binding of myosin heads to the thin filament was modeled as described previously^28^. Because the kinetics of OM binding to myosin has not been studied, we did not allow the exchange of OM with the myosin heads, but set the proportion of myosin with OM bound based the simulated concentration of OM and using a K_D_ of 100 nM for motility simulations and 1.2 μM for cardiomyocyte simulations. Measured phosphate release rates from solution were used for the attachment rates to an undecorated actin filament. Model parameters were taken from the literature when possible, with only one parameter being adjusted to fit experimental data (Supplemental Table 5). At each concentration of OM and calcium, 1000 steps of the simulation were run 1000 times. For the cardiomyocyte simulations, the actin position was held constant (to simulate isometric conditions) and the average force produced during the last half of the simulations was recorded. To simulate motility assay results, the simulations were run without any actin regulation and the total distance the actin filament traveled was divided by the simulation time was used as the measured velocity. For the OM titration curves (Figure 5c), the simulations plotted were run at pCa 6.4 since this was when the simulated force was approximately 15% activated at zero OM, as was pCa 6.0 in Nagy et al^11^. Forces for Figures 5c-d were scaled to match the data from Nagy et al by first subtracting the simulated force at 0 OM at pCa 9 (simulated passive force), then multiplying by a constant factor set to scale the force produced at pCa 4.75 (with no OM) in the simulations to that measured by Nagy et. al under the same conditions. This same constant factor (0.306) was used across all parameters, OM concentrations, and pCA values.

## ACKNOWLEDGEMENTS

This work was supported by National Institutes of Health grant R35-GM118139 to YEG, P01-GM087253 to YEG and EMO, R01-HL133863 to DAW and EMO, R00 HL123623 to MJG and a National Science Foundation Graduate Research Fellowship to M.S.W.

## AUTHOR CONTRIBUTIONS

MSW performed and analyzed optical trap experiments, stopped-flow experiments, and simulations. MJG performed optical trap experiments and made the initial observations. BB and DAW provided proteins and performed and analyzed motility assays. EMO and YEG directed research and contributed to analysis. EMO, YEG, and MSW wrote the manuscript. All authors reviewed and edited the manuscript.

## COMPETING INTERESTS

The authors declare they have no competing financial interests related to this work.

